# Immunization with peptide encapsulated within synthetic spores activates T cell responses and reduces tumor growth

**DOI:** 10.1101/2025.02.27.640614

**Authors:** Domenico D’Atri, Elena Tondini, Federico Machinandiarena, Minsuk Kong, Alilin Mia, Devorah Gallardo, Kandice Tanner, Stephen M. Hewitt, David J. Fitzgerald, Kumaran S. Ramamurthi

**Author notes:** To whom correspondence should be addressed (D.J.F.), (K.S.R.). Department of Food Science and Technology, Seoul National University of Science and Technology, Seoul 01811, South Korea (Minsuk Kong).

## Abstract

Peptide-based therapeutic immunizations represent safe approaches to elicit antigen-specific T cell responses, but their broad utility remains limited due to poor immunogenicity and short *in vivo* stability due to rapid degradation and clearance. Here we employed synthetic bacterial spore-like particles, “SSHELs”, made entirely of biocompatible materials, to deliver a model peptide antigen in the absence of additional adjuvants. SSHELs carrying the peptide antigen were internalized by dendritic cells and SSHEL-delivered peptides were then processed and cross-presented *in vitro* and *in vivo* more efficiently than free peptides. Further, SSHEL-delivered peptides elicited effective antigen-specific T cell expansion in a manner that was dependent on particle size and peptide presentation mode (encased peptides were superior to surface-attached peptides). In a mouse melanoma model expressing the antigen ovalbumin, therapeutic immunization reduced tumor size and increased survival. We propose that SSHELs are a self-adjuvanting peptide delivery system that mimics a natural presentation to elicit a robust immune response.

## INTRODUCTION

Therapeutic cancer peptide immunogens may be used to treat established cancers by provoking T cell responses against tumor antigens that might otherwise avoid detection. The use of peptide antigens in cancer immunotherapy has been widely explored due to their ability to inhibit and reduce tumor growth with relatively low toxicity and high specificity, combined with the relatively low of cost of peptide synthesis and purification (*1–3*). Peptide antigens aim to stimulate CD8^+^ cytotoxic T lymphocytes (CTLs), harnessing cellular immunity to achieve the efficient eradication of tumors (*4–7*). However, their widespread clinical use has been limited due to poor immunogenicity and low *in vivo* stability, which requires the use of adjuvants that may provoke side effects that can reduce tolerability (*8–10*). One solution to overcoming the reliance on adjuvants has been to use targeted delivery systems such as liposomes to deliver the antigen, sometimes with an encased adjuvant (*11*). Although liposomes provide several advantages such as biocompatibility, biodegradability, and high immunogenicity, liposomes may suffer from a relatively short biological half-life due to oxidation and hydrolysis, and low solubility (*12*).

We previously reported the use of synthetic bacterial spore-like particles termed “SSHELs” (<u>S</u>ynthetic <u>S</u>pore <u>H</u>usk-<u>E</u>ncased <u>L</u>ipid bilayer) (*13*), for the targeted delivery of doxorubicin to HER2-positive ovarian tumors using a murine xenograft model (*14*). Spores of the bacterium *Bacillus subtilis* are encased in a proteinaceous coat composed of ∼80 proteins (*15*). The coat is built atop two structural proteins that form a basement layer: a 26 amino-acid-long small protein named SpoVM that anchors the cytoskeletal protein SpoIVA that hydrolyzes ATP to irreversibly polymerize (*16–19*). We reconstituted the basement layer of the spore coat *in vitro* by first applying a lipid bilayer to 1 µm-diameter silica beads to generate spherical supported lipid bilayers (*20, 21*), then adsorbing synthesized SpoVM peptide to the membrane, followed by incubating purified SpoIVA in the presence of ATP, which polymerized around the SpoVM-covered spherical supported lipid bilayers to generate SSHEL particles (*22*). Previously, we demonstrated that intramuscular inoculation of mice with SSHELs covalently displaying an inactivated bacterial staphylococcal alpha toxin variant elicited the production of neutralizing antibodies and memory T cells against the antigen that resulted in improved protection against *Staphylococcus aureus* infection in a murine bacteremia model (*23*). Here, we investigated the self-adjuvanting properties of SSHELs in delivering a peptide that harbors the “SIINFEKL” amino acid sequence derived from ovalbumin (*24*). We report that SSHELs carrying the model peptide are internalized by dendritic cells and that the core SIINFEKL antigen is processed and cross presented in the context of MHC-I molecules *in vitro* and *in vivo*. Furthermore, we observed that particle size as well as mode of delivery (covalently surface displayed peptide or peptide encased within the particle) both influence the efficacy of cross presentation by dendritic cells. Intramuscular inoculation with SSHELs promoted peptide-specific CD8+ T cell activation and expansion with robust cytokine release. Finally, using a syngeneic murine melanoma tumor model together with a cancer cell line that expresses ovalbumin, we observed that inoculation with SSHELs delivering the model peptide reduced tumor burden and increased survival. We propose that SSHELs represent a versatile peptide antigen delivery system that can be easily adjusted to suit specific therapeutic requirements.

## RESULTS

### SSHEL size and mode of peptide presentation affect cross presentation efficiency

As a model antigen, we chose the well-characterized peptide sequence SIINFEKL, which is derived from ovalbumin, is efficiently displayed in the context of an MHC-I molecule, and for which multiple reagents exist to detect its cross-presentation (*25*). However, instead of using the bare SIINFEKL epitope, we employed a longer peptide derived from ovalbumin (Fig. 1A) in which the SIINFEKL sequence was embedded, to ensure that the peptide was internalized and processed before it was cross presented, and to increase cross-presentation efficiency (*26, 27*). Next, to understand the best method of using SSHELs to deliver the epitope, we sought to test relative efficiencies of SIINFEKL cross-presentation to 1) determine if the peptide could be delivered by chemical crosslinking on the SSHEL surface (via cross-linking to SpoIVA) or by encapsulation within the framework of the particles, and 2) to monitor response based on the size of the SSHEL particle (Fig. 1B). We therefore used SSHEL particles that were built atop mesoporous silica beads (which allows for the encapsulation of cargo) that were either 500 nm or 1 µm in diameter. We then either encapsulated an equivalent amount of peptide inside the silica constructing the SSHEL particle or crosslinked an equivalent amount of peptide to the surface of the particle using a unique Cys residue engineered into SpoIVA and a unique Cys residue engineered to the carboxy terminus of the peptide. The four resulting SSHEL particles were then incubated with immortalized dendritic cells (MutuDC1940) and cross-presentation was measured by flow cytometry using a fluorescently labeled antibody that specifically recognizes the SIINFEKL peptide in the context of MHC-I. Displaying the antigen on the SSHEL surface resulted in ∼1.5-fold higher cross-presentation when delivered by 500 nm SSHELs compared to 1 µm SSHELs. (Fig. 1C). In contrast, delivering the peptide alone with no association to SSHELs did not result in cross-presentation that was appreciably above background fluorescence. When the antigen was encapsulated, 500 nm SSHELs showed a ∼7-fold increase in SIINFEKL cross-presentation compared to 1 µm SSHELs (Fig. 1D). Thus, not only particle size, but method of antigen delivery evidently influences the efficiency of antigen cross-presentation. We therefore chose 500 nm particles with encapsulated peptide to deliver the SIINFEKL epitope for all subsequent experiments (hereafter, “SSHEL^EncPep^”). To test if the SSHEL platform specifically contributes to efficient peptide delivery, we compared cross-presentation of the epitope to peptide delivered by liposomes made from phosphatidylcholine (the same lipids used to construct SSHELs). The results in Fig. 1E indicate that SSHEL^EncPep^ resulted in a ∼700-fold higher SIINFEKL cross-presentation compared to peptide encased in liposomes. The data therefore suggest that 500 nm SSHELs that deliver the model antigen via encapsulation result in optimal cross-presentation of the epitope in cultured dendritic cells.

**Figure 1.**
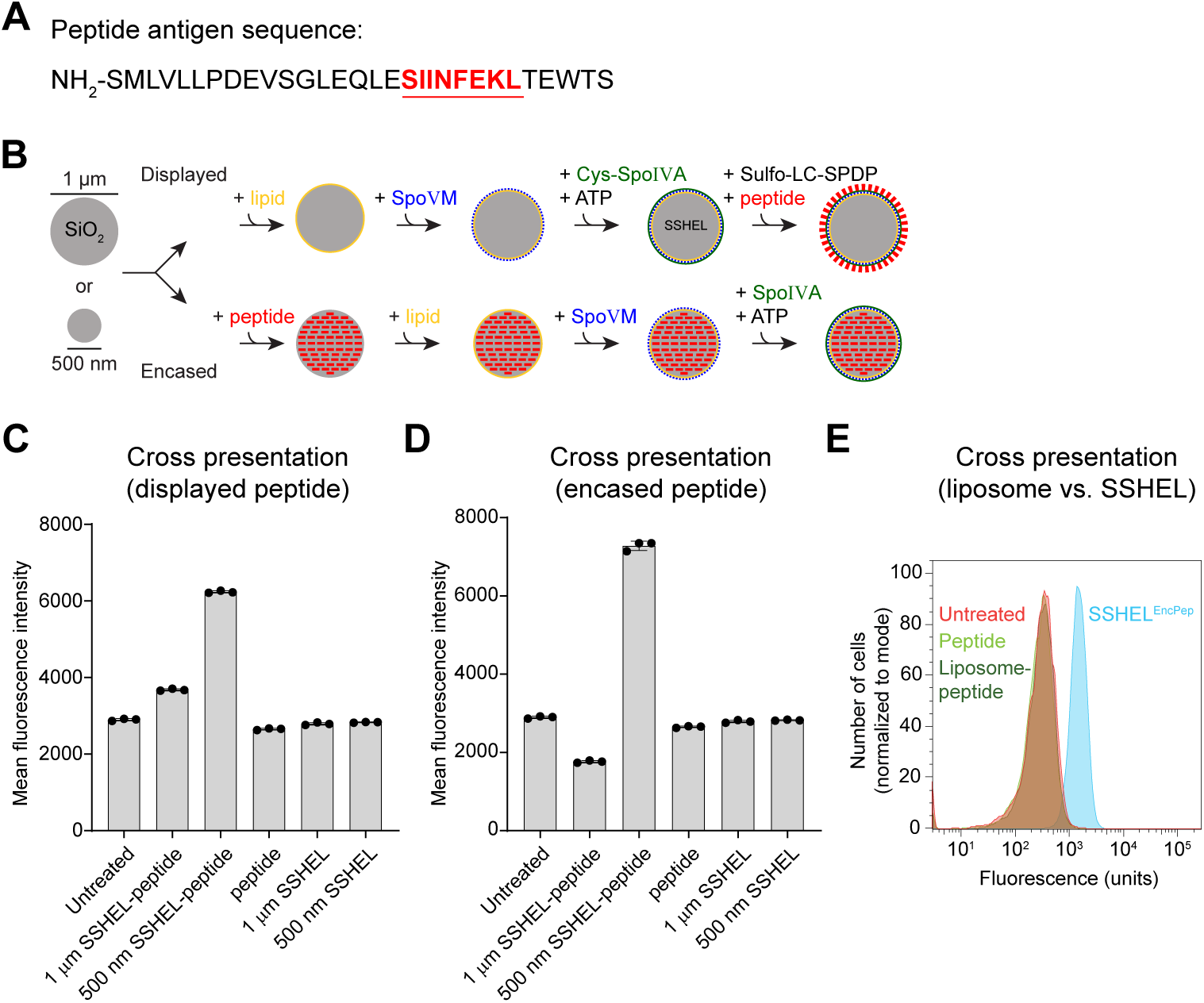
SSHEL-mediated peptide antigen delivery leads to cross presentation of the antigen *in vitro*. (A) Schematic of SSHEL assembly. 1 µm- or 500 nm-diameter mesoporous silica beads (gray) were (top) covered with a lipid bilayer (yellow), then SpoVM peptide (blue), then polymerized SpoIVA protein (green). Free amines on peptide (red) were covalently linked to a single engineered Cys in SpoIVA via Sulfo-LC-SPDP cross linker. Alternatively, mesoporous silica beads were (bottom) loaded with peptide, then covered with a lipid bilayer (yellow), then SpoVM peptide (blue), then polymerized SpoIVA protein (green). (B) Ovalbumin-derived peptide sequence harboring the SIINFEKL epitope (red, underlined). (C-D) Cross-presentation of SIINFEKL epitope by immortalized dendritic cells measured using flow cytometry when incubated with SSHELs of indicated size that (C) displayed or (D) encased the peptide. Bars indicate mean; errors: S.D. Data points represent independent replicates. (E) Cross-presentation of SIINFEKL epitope by immortalized dendritic cells when incubated with peptide encased in (blue) 500 nm SSHELs, (dark green) liposomes, or (light green) free peptide.

### Cross-presentation of the epitope requires particle uptake and peptide processing

We next tested if the cross-presentation of the SIINFEKL epitope was due to SSHEL^EncPep^ internalization, followed by processing of the encapsulated longer peptide to display the mature antigen (*28*). Previously, we demonstrated that SSHELs bound to a surface receptor (HER2) on an ovarian cell line can be actively internalized via a macropinocytosis-like process that involves actin polymerization. To test if the cultured dendritic cells also require particle uptake to cross-present the epitope, we measured cross-presentation *in vitro* in the presence of the actin polymerization inhibitor, latrunculin A. Addition of latrunculin A reduced peak cross-presentation compared to absence of the drug (Fig. 2A); as controls, delivery of the antigen via liposome (Fig. 2B) or free peptide (Fig. 2C) failed to result in cross-presentation in the presence or absence of latrunculin A. Finally, varying the concentration of encapsulated peptide that was delivered by SSHELs to dendritic cells *in vitro* revealed that increasing peptide concentrations resulted in increased cross-presentation (Fig. 2D-E), but delivering free peptide did not result in increased cross-presentation, indicating that the full-length peptide must be processed before the SIINFEKL epitope may be cross-presented. Taken together, the data are consistent with a model in which SSHEL^EncPep^ is internalized, the encased peptide is released and processed, and the internal SIINFEKL peptide sequence is subsequently cross-presented.

**Figure 2.**
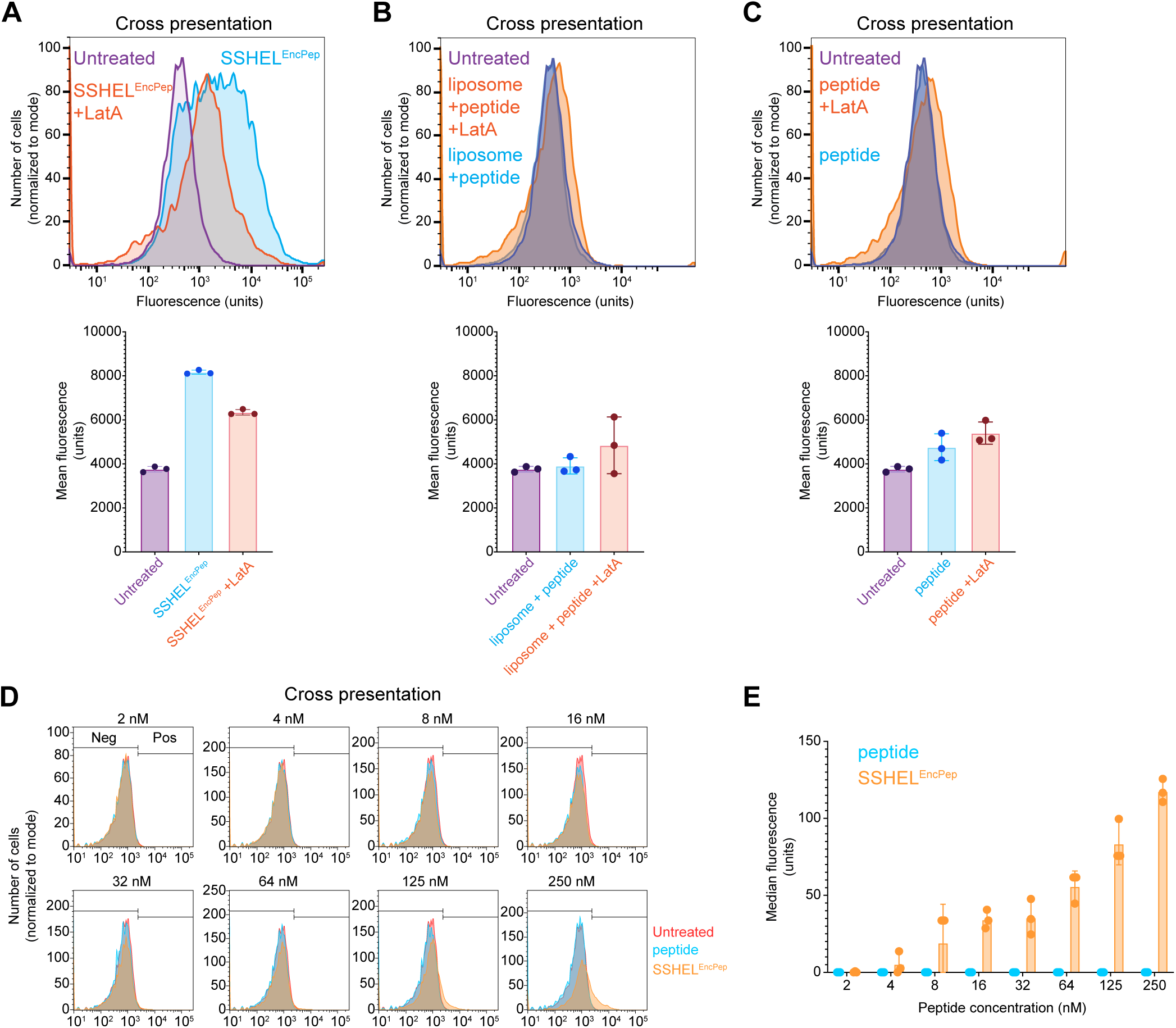
Cross presentation of the SIINFEKL epitope requires particle uptake and peptide processing. (A-C) Cross-presentation of SIINFEKL epitope by immortalized dendritic cells measured using flow cytometry when incubated in the absence (purple) or presence of (A) SSHEL^EncPep^, (B) liposomal-encased peptide, or (C) peptide alone (blue), in the additional presence of latrunculin A (orange). Fluorescence intensities derived from each histogram are shown below. Bars indicate mean; errors: S.D. (D) Cross-presentation of SIINFEKL epitope by immortalized dendritic cells when incubated with indicated concentration of peptide as (blue) free peptide or (orange) SSHEL^EncPep^. (E) Fluorescence intensities derived from histograms in (D). Bars: median fluorescence; errors: S.D. All data points represent independent replicates.

### SSHEL-delivered antigen is efficiently cross-presented in vivo

Next, we compared how freshly isolated mouse dendritic cells cross-present the SSHEL-delivered SIINFEKL epitope and if SSHEL^EncPep^ would elicit the release of cytokines. Dendritic cells were isolated from freshly harvested spleens from C57Bl/6 immunocompetent mice and incubated with free full-length peptide harboring the SIINFEKL epitope, or 1 µm- or 500 nm-diameter SSHELs either displaying or encasing the peptide, after which cross-presentation of the SIINFEKL epitope was assessed using flow cytometry. Similar to cultured dendritic cells, delivery via 500 nm SSHEL^EncPep^ resulted in the most efficient cross-presentation of the SIINFEKL epitope (Fig. 3A, red histogram). In contrast, though, incubation of primary dendritic cells with free peptide also resulted in efficient cross-presentation of the epitope (Fig. 3A, orange histogram).

**Figure 3.**
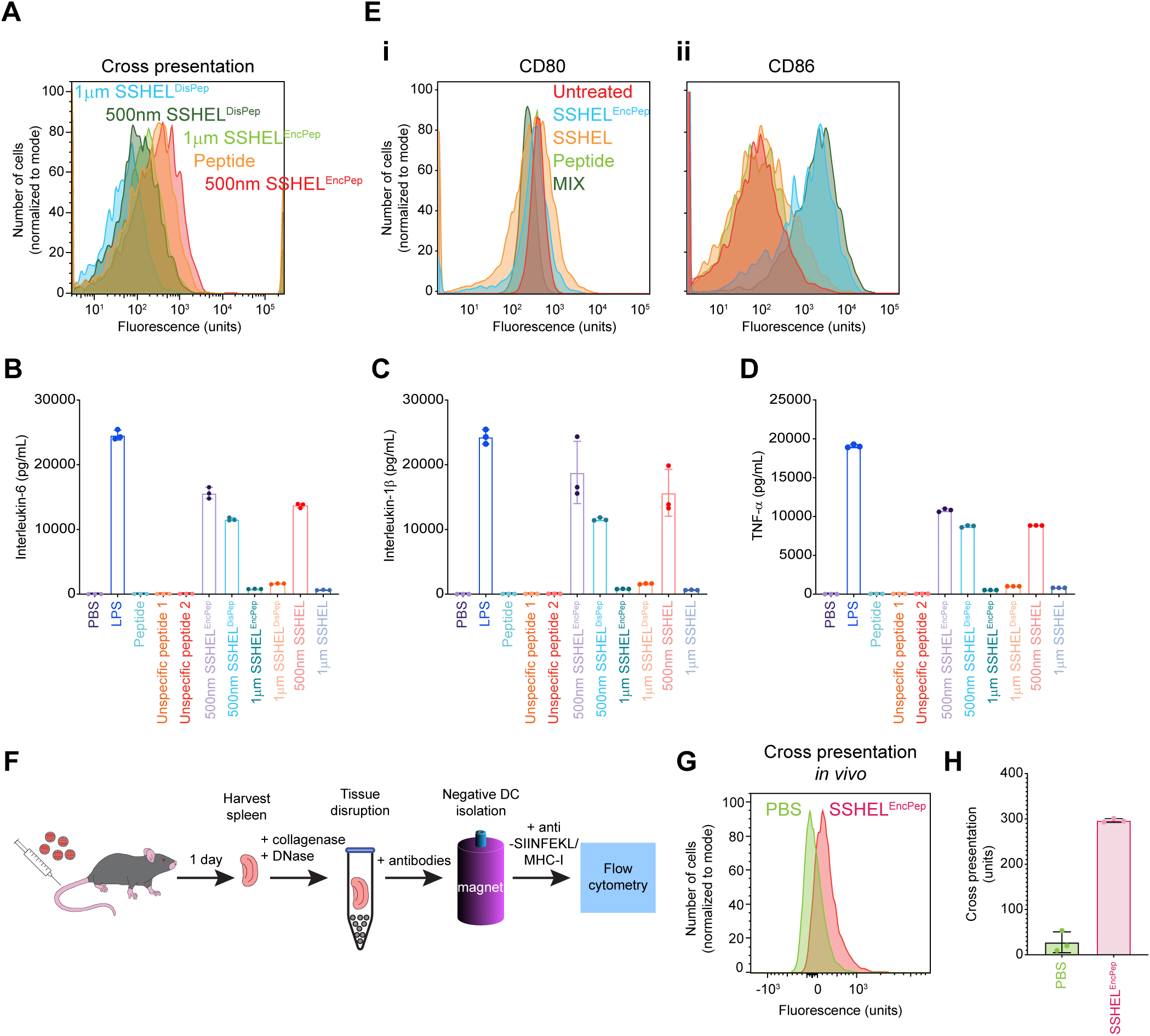
Cross presentation of SSHEL-delivered SIINFEKL epitope *in vivo*. (A) Cross-presentation of SIINFEKL epitope by freshly isolated mouse dendritic cells using flow cytometry when incubated with (orange) peptide alone, (blue) 1 µm or (dark green) 500 nm SSHELs displaying the peptide, or (light green) 1 µm or (red) 500 nm SSHELs encasing the peptide. (B-D) Release of cytokines by freshly isolated mouse dendritic cells after incubation with the indicated antigen. LPS: lipopolysaccharide; unspecific peptide 1-2: two different peptides in which the sequence of the SIINFEKL-containing peptide was scrambled (see Experimental Procedures). (E) Schematic representation of immunization protocol. C57Bl/6 mice were intravenously injected with 500 nm SSHEL^EncPep^. After one day, dendritic cells were isolated from harvested spleens stained and SIINFEKL cross presentation was detected using flow cytometry. (F-G) Cross presentation of SIINFEKL peptide by dendritic cells isolated from mice inoculated with PBS (green) or SSHEL^EncPep^.

We then examined the culture medium for cytokines that were released by the dendritic cells after incubation with the various antigens. Incubation with 500 nm SSHELs, but not 1 µm SSHELs (without any peptide), was sufficient to trigger release of IL-6, IL-1β, and TNF-α (Fig. 3B-D, light red and light blue), which are cytokines required to stimulate CD8+ cytotoxic T cells. Accordingly, 500 nm SSHEL^EncPep^ and SSHELs harboring surface-displayed peptide (SSHEL^DisPep^) more robustly triggered release of these cytokines compared to the corresponding 1 µm SSHELs (Fig. 3B-D, light purple and dark green). In contrast, although incubation with free peptide promoted efficient cross-presentation of the SIINFEKL epitope by freshly isolated dendritic cells (Fig. 1A), it did not trigger the robust cytokine release (Fig. 1B-D, light green), similar to the activities elicited by two different unspecific peptides (Fig. 1B-D, dark orange and dark red). Examining these dendritic cells by flow cytometry revealed the slight overproduction of CD80 and the higher overproduction of CD86, two receptors that are required for T cell activation, when the dendritic cells were incubated with SSHEL^EncPep^ (Fig. 3Ei-ii).

To test if epitope delivery via SSHELs results in successful cross-presentation *in vivo*, we intravenously injected C57Bl/6 mice with SSHEL^EncPep^, harvested the spleens after 1 day, and measured the cross-presentation of the SIINFEKL epitope by isolated dendritic cells using flow cytometry (Fig. 3F). Compared to dendritic cells isolated from mice injected with PBS, injection with SSHEL^EncPep^ resulted in a ∼10-fold increase in cross-presentation signal (Fig. 3G-H) in the context of an intact immune system. We therefore conclude that delivering enclosed peptide via 500 nm SSHELs results in cross-presentation of the processed antigen *in vivo* and that this mode of delivery likely activates the dendritic cell to prime cytotoxic T cells.

### Inoculation with SSHEL^EncPep^ promotes effective T cell expansion

To validate whether the SSHEL^EncPep^ is a suitable platform for delivering antigens, we performed inoculations on C57Bl/6 mice via two different routes: intravenous (since some antigens have reported a better response to IV injection (*29, 30*)) and intramuscular, both without using adjuvants, and monitored the priming of cytotoxic T cells. Mice were injected twice over two weeks, peripheral blood was collected, and CD3+/CD8+ double positive T cells were isolated and analyzed by flow cytometry for their ability to bind to MHC-I/SIINFEKL (Fig. 4A; Fig. S1). Mice inoculated intravenously with SSHEL^EncPep^ displayed a ∼3-fold increase in SIINFEKL-specific cytotoxic T cells compared to mice injected with free peptide (Fig. 4C). However, inoculating mice intramuscularly with SSHEL^EncPep^ resulted in ∼60-fold increase in SIINFEKL-specific cytotoxic T cells compared to mice inoculated with free peptide (Fig. 4B-C), and ∼6-fold increase compared to IV injection with SSHEL^EncPep^ (Fig. 4C).

**Figure 4.**
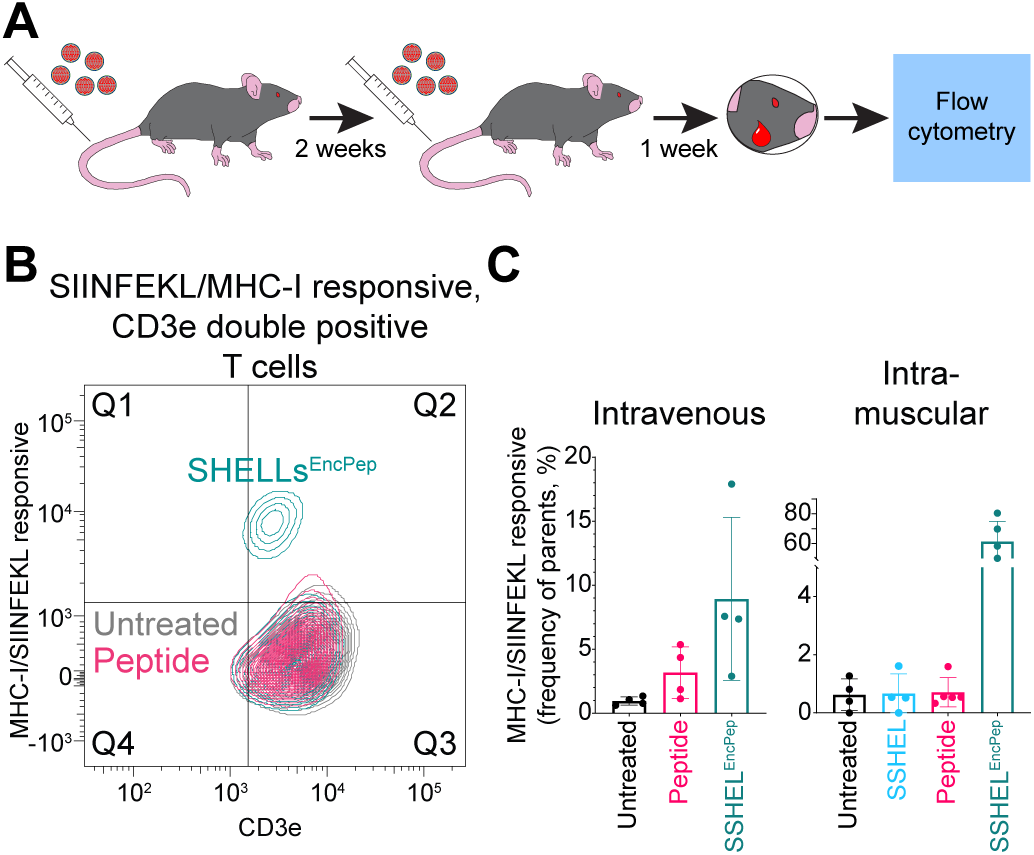
Intramuscular inoculation with SSHEL^EncPep^ promotes effective T cell expansion against SIINFEKL. (A) Schematic representation of inoculation protocol. C57Bl/6 mice were injected intravenously or intramuscularly with PBS, peptide, 500 nm SSHEL, or 500 nm SSHEL^EncPep^ and boosted via the same inoculation route 2 weeks later. After 1 week, blood sample was collected, CD3e/CD8a double positive cytotoxic T cells were isolated by flow cytometry and analyzed for their ability to bind to MHC-I/SIINFEKL. (B) Representative flow cytometry plot of isolated CD3e/CD8a positive from mice inoculated intramuscularly with 500 nm SSHEL^EncPep^ that recognize the SIINFEKL tetramer. (C) Percentage of CD3e/CD8a positive cytotoxic T cells that recognize SIINFEKL when presented in MHC-I harvested from mice injected (left) intravenously or (right) intramuscularly with (black) PBS, (pink) free peptide, (blue) SSHEL, or (green) SSHEL^EncPep^.

We then compared CD8+ T cell activation and cytokine release after inoculation via both routes by harvesting spleens from mice that were inoculated as described above, isolating splenocytes, activating them *ex vivo* with dendritic cells displaying the SIINFEKL epitope, and measuring cytokine production by staining with specific antibodies followed by flow cytometry. Mice inoculated intravenously with SSHEL^EncPep^ induced an ∼2-fold higher overall cytokine response compared to mice that were inoculated with peptide only (Fig. 5A; Fig. S2). Examining the polyfunctionality of the cytokine response, intravenous injection of either peptide alone or SSHEL^EncPep^ resulted in an IFNγ-skewed response (Fig. 5B). In contrast, intramuscular inoculation with either free peptide or SSHEL^EncPep^ resulted in higher overall activation of SIINFEKL-specific CD8+ T cells, with SSHEL^EncPep^ inoculation triggering ∼50% higher overall cytokine response compared to free peptide (Fig. 5C). More importantly, though, examining the distribution of the released cytokines revealed that the fraction of CD8+ T cells producing multiple cytokines was increased when the antigen was delivered intramuscularly by SSHEL^EncPep^, with 18% of CD8+ T cells producing IFN-γ, IL-2, and TNF-α, compared to just 0.9% for intramuscular inoculation with peptide alone (Fig. 5D). Together, the results indicate that intramuscular injection of peptide antigen encapsulated in 500 nm SSHELs, even without an adjuvant, enhances the activation of antigen-specific cytotoxic T cells that display a polyfunctional cytokine profile.

**Figure 5.**
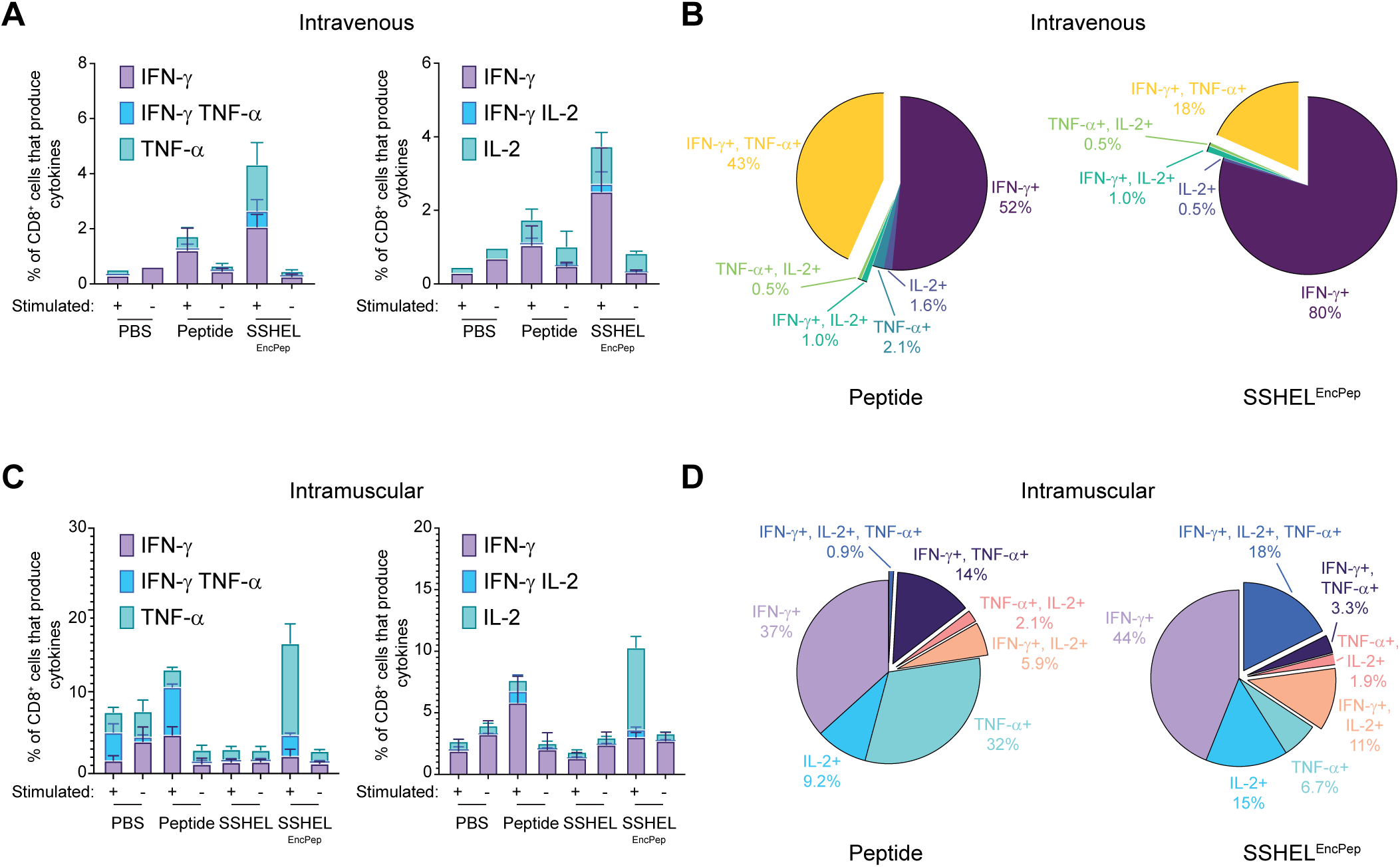
Intramuscular inoculation with SSHEL^EncPep^ promotes robust T cell activation and cytokine release *in vivo*. (A-D) Cytokine release was assayed from cytotoxic T cells isolated from spleens harvested from C57Bl/6 mice injected (A-B) intravenously or (C-D) intramuscularly with PBS, peptide, or 500 nm SSHEL^EncPep^ twice (inoculation schedule shown in Fig. 4A) one week after second injection. Isolated splenocytes were stimulated by incubation with MutuDC1940 cells presenting SIINFEKL antigen, and T cell activation and cytokine release was measured using flow cytometry. (A, C) Fraction of total CD8 positive T cells releasing (left) IFN-γ, TNF-α, or both, or (right) IFN-γ, IL-2, or both. (B, D) Fraction of total CD8 positive T cells harvested from mice inoculated with (left) peptide or (right) SSHEL^EncPep^ releasing the indicated cytokine(s) after restimulation.

### Therapeutic immunization with SSHEL^EncPep^ increases survival in a mouse melanoma model

To test the efficacy of inoculation with SSHELs delivering the SIINFEKL antigen, we chose a syngeneic tumor model using a mouse melanoma cell line that expresses ovalbumin (B16-N4-OVA). In trial 1, a subcutaneous tumor was first established. When tumor size was ∼100 mm^3^, the first inoculation was administered, followed by a second inoculation 7 days later. Whereas the mean tumor size in PBS-treated mice increased to ∼950 mm^3^ in 13 days (n = 8 mice), the mean tumor size in mice inoculated with SSHEL^EncPep^ reached only ∼450 mm^3^ (Fig. 6A; individual tumor sizes reported in Fig. S3A; mean mouse weights reported in Fig. S4A). Additionally, individual tumor sizes in mice in the SSHEL^EncPep^ treatment group that survived beyond day 18 largely plateaued, compared to mean tumor sizes in the other treatment groups (Fig. S3A). As controls, inoculation with peptide alone or SSHEL alone resulted in tumor growth rates that were similar to treatment with PBS (Fig. 6A). Consistent with this pattern, mice inoculated with SSHEL^EncPep^ also exhibited increased survival in this trial (Fig. 6B). In trial 2, we tested two more control groups: mice inoculated with SSHELs harboring an unspecific peptide that is also cross-presented in MHC-I (*31*), or mice co-inoculated with empty SSHELs alongside (external) SIINFEKL-harboring peptide. In this trial, inoculation with SSHEL^EncPep^ initially resulted in reduced mean tumor growth compared to all treatments except for inoculation with free peptide (Fig. 6C). However, individual tumors in the free peptide inoculation group nonetheless reached the size at which the mouse needed to be sacrificed by day 25, whereas the growth of tumors in mice in the SSHEL^EncPep^ inoculation group largely plateaued (Fig. S3B). As a result, consistent with the pattern in tumor sizes, inoculation with SSHEL^EncPep^ resulted in increased survival compared to all other treatment groups (1.3-fold longer than the PBS treated group, and 1.25-fold longer than mice inoculated with peptide only; Fig. 6D). We also conducted a third trial, consisting of just 4 mice per treatment group, to assess the pathology of the tumors. In this trial, all mice were sacrificed at day 15, when the first mouse reached a protocol endpoint (tumor size). Tumors were resected, measured, and submitted for histological analysis. As observed in the other trials, inoculation with SSHEL^EncPep^ resulted in reduced mean tumor sizes (Fig. S5 A-B). Hematoxylin and eosin-stained sections of all four groups were reviewed blind, and all sections demonstrated a substantial infiltrate of melanophages (macrophage with phagocytized melanin), zones of necrosis and variable inflammatory response. Tumor size was the only feature that distinguished the groups on pathology review. However, distinct pathologic features that allowed distinguishing differences between the groups included percent necrosis (indicating rapid tumor growth), which was higher in the untreated group (60%), compared to 10%-40% for the free peptide group, 20%-40% for mice inoculated with SSHELS only, and 20-40% for the SSHEL^EncPep^. In sum, we conclude that therapeutic immunization with a peptide antigen delivered by SSHELs increases survival and reduces tumor burden in a syngeneic mouse tumor model.

**Figure 6.**
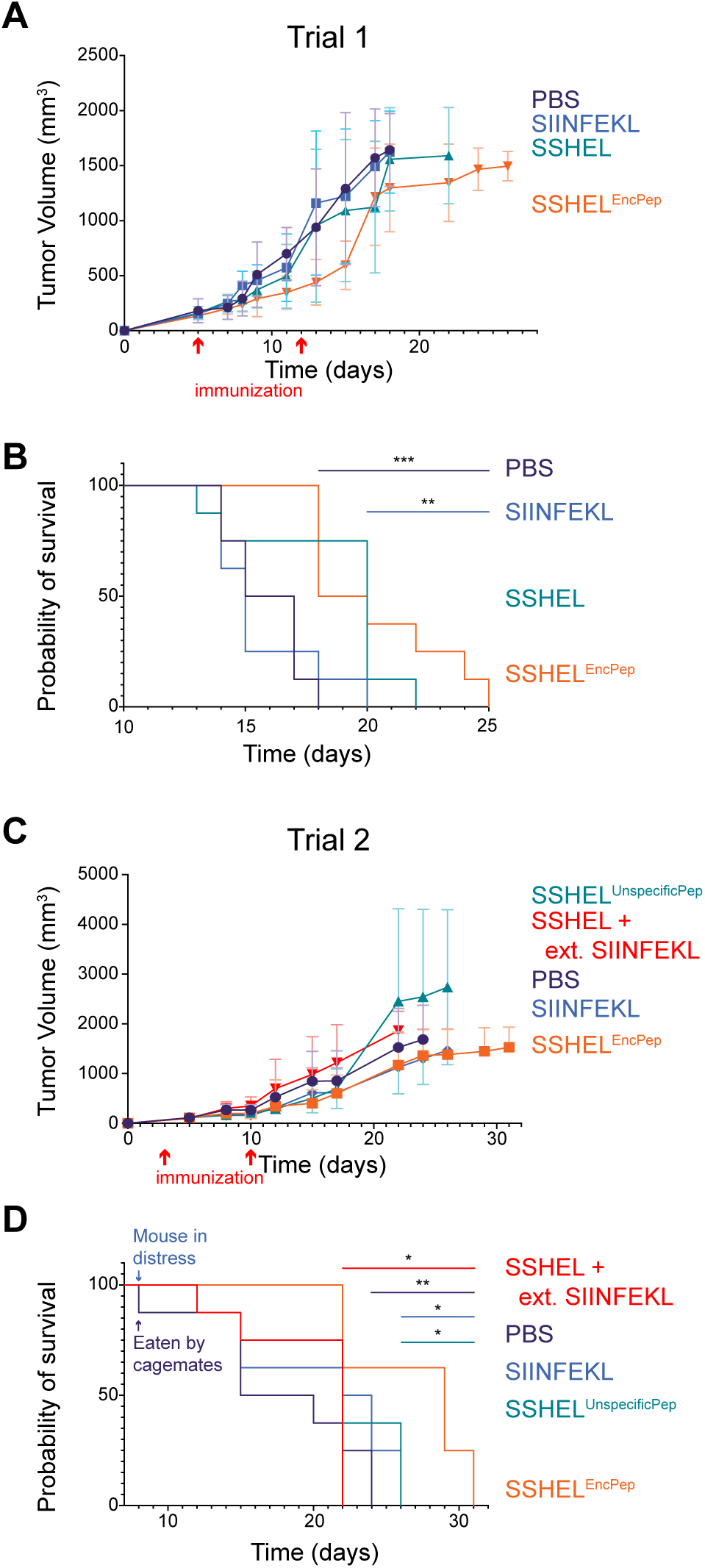
Immunization with SSHEL^EncPep^ reduces tumor size and increases survival. (A-D) Subcutaneous B16-N4-OVA tumor (mouse melanoma expressing ovalbumin) was introduced in C57Bl/6 mice (8 mice/treatment group). When tumors reached ∼100 mm^3^, the first inoculation was administered (7 µg peptide), followed by a second inoculation two weeks later. (A, C) Tumor size and (B, D) survival. (A-B) In trial 1, mice were treated with PBS (purple), SIINFEKL peptide alone (blue), SSHEL alone (green), or SSHEL^EncPep^ (orange). (C-D) In trial 2, mice were treated with PBS (purple), SSHEL^UnspecificPep^ (green), SSHEL external SIINFEKL peptide (red), SIINFEKL peptide alone (blue), or SSHEL^EncPep^ (orange).

## DISCUSSION

In this study, we assessed the efficacy of SSHELs as a delivery system for therapeutic peptide antigens. We observed that peptide antigens delivered by SSHELs were processed, cross-presented by dendritic cells, and elicited antigen-specific T cell expansion. Interestingly, SSHELs present a rare opportunity to vary not only particle size, but also mode of delivery (surface display via covalent conjugation or encapsulation) without altering the chemical makeup of the particle. Previous studies have suggested that particle shape and size can affect transport to the lymph nodes and therefore pharmacodynamics (*32, 33*). Other studies reported that varying particle size may elicit different types of immune responses. For example, delivering full length ovalbumin conjugated to differently sized carboxylated polystyrene microspheres co-injected with adjuvants revealed that virus-sized particles (40-50 nm) elicited robust combined cellular and humoral responses (*34*). Another study reported that slightly larger particles engineered from lecithin/glyceryl monostearate polymers induced strong antibody and cellular immune responses at the 230 nm size range (*35*). Here, we report additionally that 500 nm SSHELs that encased the peptide, rather than those displaying the peptide, maximized cross-presentation and antigen-specific T cell expansion. Since the SSHEL surface is readily modifiable, it will be interesting to test if targeting moieties attached the surface of SSHELs that bind directly to antigen presenting cells may further increase the efficiency of antigen delivery.

SSHELs therefore represent a versatile peptide-based antigen delivery system which combines the ability of delivering peptides (theoretically a single species or a combination of multiple peptides containing multiple epitopes) with its own auto-adjuvant properties for therapeutic cancer immunization. Indeed, we have previously observed that immunizing with SSHELs generates antibodies against the SpoIVA protein, but not a memory T cell response, suggesting that repeated inoculation using SSHELs with different antigens conjugated to SpoIVA may be tolerated (*23*). Finally, physical modification of SSHELs (for example, size and delivery method) may further potentiate dendritic cell functions and effector T cell activation and expansion, which may translate into more effective anti-tumor immunity *in vivo*.

## MATERIALS AND METHODS

### SSHEL assembly

Synthesized ovalbumin peptide containing the SIINFEKL epitope (sequence SMLVLLPDEVSGLEQLE<u>SIINFEKL</u>TEWTS; Genscript, 95% - 99% purity) was dissolved in DMSO to a final concentration of 10 mM. For SSHELs that encapsulated the antigen, 3 mg of silica beads (1 µm or 0.5 µm diameter, 10 nm pores, amine-functionalized surface) were incubated overnight with a 0.2 mM solution of peptide at room temperature in a dark environment. After encapsulation, beads were harvested by centrifugation at 13000 × *g* for 5 min, and the unencapsulated peptide in the supernatant was quantified using HPLC (by comparing the integrated area under the elution curve compared to a standard curve constructed using known concentrations of peptide) to determine the cargo load. After encapsulation, SSHELs were built largely as described in previously (*22*). Briefly, ∼200 µl (∼10 mg) peptide-loaded beads were incubated with ∼5 mg 1,2-dioleoyl-sn-glycero-3-phosphocholine (DOPC) liposomes in PBS for 1 h at room temperature with gentle inversion. Next, lipid coated beads were incubated with 10 μM synthesized SpoVM peptide (Biomatik) and 1.5 μM purified His_6_-SpoIVA in 50 mM Tris at pH 7.5, 4 mM ATP, 10 mM MgCl_2_, 400 mM NaCl overnight at room temperature with gentle inversion. His_6_-SpoIVA was produced and purified from *E. coli* “ClearColi” BL21(DE3) pJP120, which expresses a modified LPS that does not trigger endotoxic response, as described previously (*14, 36*). For SSHELs displaying the peptide antigen, the peptide was first cross-linked to SpoIVA using Sulfo-LC-SPDP (ThermoFischer, Cat No. 21650) according to the manufacturer’s instructions. To perform this cross-linking reaction, a Cys-less SpoIVA variant was used. After cross-linking the antigen to SpoIVA, SSHELs were assembled as described above.

### Cell culture

MutuDC1940 immortalized mouse dendritic cells (Applied Biological Materials Inc.) were cultured at 37 °C, 5% CO₂ in IMDM Glutamax medium (ThermoFischer Scientific Cat No. 31980030) supplemented with heat-inactivated 10% FBS (ThermoFischer Scientific Cat No. A5256801), 50 µM β-mercaptoethanol (Gibco Cat No. 21985-023), 10 mM HEPES (Corning Cat No. 25-060-CI), 1% Penicillin/Streptomycin Solution (Sigma Aldrich Cat No. P0781).

### In vitro antigen detection

At day -1 50,000 MutuDC1940 cells per well were plated in a 96 well plate and cultured overnight. At day 0 the cells were stimulated with the peptide reporter antigen (2 to 250 nM) free or delivered by SSHELs and incubated overnight. At day 1 cells were washed three times with PBS and then gently detached using a 0.1% Trypsin-EDTA solution. The cells were then incubated for 30 min at room temperature with antibody against H-2Kb bound to SIINFEKL (Biolegend, Cat No. 141604), and then washed three times using flow cytometry buffer (PBS, BSA 1%, NaN_3_ 0.1%). Cross-presentation was evaluated by flow cytometry using BD FACSymphony A5 (BD Biosciences). All data were analyzed using FlowJo 10.10.0 (Becton Dickinson & Company).

### Ex vivo antigen detection

Spleens from C57Bl/6 immunocompetent mice were isolated and incubated with Collagenase IV (250 U ml^-1^; Merck, Cat. No. 11088866001) and DNAse (50 µg ml^-1^) (Merck, Cat. No. 10104159001) at 37 °C. After degradation, a single cell suspension was prepared and enriched for dendritic cells using EasySep Mouse Pan-DC Enrichment Kit (STEMCELL Technologies, Cat No. 19763). After incubation, cells were co-incubated with the reporter antigen (free or delivered by SSHELs) and cultured for 6 h at 37 °C, 5% CO₂ in IMDM Glutamax medium (ThermoFischer Scientific, Cat No. 31980030) supplemented with heat-inactivated 10% FBS (ThermoFischer Scientific, Cat No. A5256801), 50 µM β-mercaptoethanol, 10 mM HEPES (Corning, Cat No. 25-060-CI), 1% Penicillin/Streptomycin Solution (Sigma Aldrich, Cat No. P0781). Cells were then collected, stained, and analyzed for H-2Kb bound to SIINFEKL using flow cytometry as described above.

In a separate experiment primary dendritic cells were isolated as described above and stimulated with SSHELs^EncPEP^ or LPS (ThermoFischer, Cat No. 00-4976-93) for 12 h. Unstimulated cells were used as a control. After stimulation, the medium was collected and analyzed for IL6, IL1β, and TNFα release using commercial ELISA kits (R&D; IL6, Cat No. 406; IL1β, Cat No. DY401; TNFα, Cat No. DY410) according to manufacturer protocol. Results were collected using a BioTek Synergy H1 spectrophotometer (Agilent). All data were analyzed using GraphPad Prism version 10.1.1 for MacOS. In a parallel experiment, the cells were stained using antibodies against CD80 (Biolegend, Cat No. 104725), CD86 (Biolegend, Cat No. 105012), CD40 (BioLegend, Cat No. 124632), or H-2Kb (Biolegend, Cat No. 116507) H-2Kb bound to SIINFEKL (Biolegend, Cat No. 141605). Marker expression was evaluated by flow cytometry using a BD FACSymphony A5 (BD Biosciences). All data were analyzed using FlowJo 10.10.0 (Becton Dickinson & Company).

### Mouse immunization and analysis of antigen-specific T cell responses

All animal procedures reported in this study that were performed by NCI-CCR affiliated staff were approved by the NCI Animal Care and Use Committee (ACUC) and in accordance with federal regulatory requirements and standards. All components of the intramural NIH ACU program are accredited by AAALAC International.

Female C57Bl/6 mice (Jackson Laboratories), aged 5-7 weeks, were inoculated (no tumor present) intravenously or intramuscularly. Seven days after the first injection the mice were boosted with a second shot following the same route. One week after the second inoculation, cheek blood samples were collected (100 to 200 μL) in 1.5 mL tube pre-filled with 20 U/tube of heparin (Millipore Sigma Cat No. H3393-250KU) to prevent blood clotting. To remove erythrocytes, the blood samples were added 1500 μL of ACK Lysing Buffer (Lonza Cat No. 10-548E) and incubated at RT for 15 minutes then the buffer action was stopped with 1 mL of PBS. The cells were centrifuged at 500 × *g*, at 4 °C for 5 min. If needed the lysis process was repeated, as suggested by the manufacturer’s protocol. Next, samples were stained using 10 µL of MHC Class I Murine iTAg Tetramer/APC – H-2 Kb OVA (SIINFEKL) (MBL International Corporation, Cat No. TB-5001-2) for 30 min at room temperature, protected from light, then washed according to manufacturer protocol. Samples were then stained with Zombie Aqua Fixable Viability Kit (BioLegend, Cat No. 423101) for live and dead staining according to manufacturer protocol. Subsequently, the cells were incubated with antibodies against CD3 and CD8 (Biolegend, Cat No. 152304 and 100734) for 30 min at 4 °C. Cells were then washed three times with flow cytometry buffer. Analysis of CD3/CD8 positive specific T cell response versus SIINFEKL was evaluated using BD FACSymphony A5 (BD Biosciences) and analyzed using FlowJo 10.10.0.

In separate experiments, we analyzed cytokine release by CD8+ T cells when presented with SIINFEKL antigen. To this end, we inoculated mice as described above and 7 days after the second injection, we harvested the spleen. Splenocytes were then isolated by crushing the spleens manually to prepare single cell suspensions. Two days prior, we seeded 50,000 MutuDC1940 cells per well in a 96 well plate. The MutuDC1940 were then stimulated the next day with 2.5 µg ml^-1^ of peptide antigen. Half of the wells were not stimulated and served as a control. On the day that the splenocytes were isolated, we co-incubated the splenocytes together with the MutuDC1940 for 6 h, then washed the cells with flow cytometry buffer thrice, then stained the with Zombie Aqua Fixable Viability Kit for live and dead staining according to manufacturer protocol. Cells were subsequently stained using antibodies against CD3 and CD8 as described above. After washing thrice with flow cytometry buffer, the cells were fixed, permeabilized and washed using Cytofix/Cytoperm (BD Industries, Cat No. 554714) according to manufacturer protocol. After fixing we stained the cells using antibodies against IL-2 (Biolegend Cat, No. 503810), TNF-α (Biolegend, Cat No. 506346), and IFN-γ (BD Pharmigen, Cat No. 557649). After washing thrice with flow cytometry buffer, the presence of internal cytokines in CD3/CD8+ T cells were evaluated using BD FACSymphony A5 and analyzed using FlowJo 10.10.0 as described above.

### Tumor trials

8 Female C57Bl/6 mice aged 5-7 weeks (Jackson Laboratories) were used in each treatment group. Mice were inoculated subcutaneously in the right flank with 5 × 10^5^ B16-N4-OVA melanoma cells. When tumor sizes reached ∼100 mm^3^ (typically 3-5 days), mice were inoculated intramuscularly using the left rear leg muscle. 7 days after the first injection the mice were inoculated again. Weight and tumor sizes were measured every two days; mice were sacrificed when tumor size exceeded 1400 mm^3^, weight loss exceeded 20%, or if mice exhibited general distress. In trial 2, the unspecific peptide used was derived from Human Papillomavirus type 18 E6 and E7 oncogenes (sequence GQAEDRAHYNIVTFCCKCDSTLRLCVK) (*31*).

For Nanostring analysis, 1-2 days after the second injection the mice were bled from the cheek. The blood (100 to 200 μL) was collected in 1.5 ml tubes pre-filled with 20 U/tube of heparin (Millipore Sigma Cat No. H3393-250KU) to prevent blood clotting. To remove erythrocytes, we added 1500 µL of ACK Lysing Buffer (Lonza, Cat No. 10-548E) and incubated at room temperature for 15 min, after which the reaction was quenched with 1 ml of PBS. The cells were then removed by centrifugation at 500 × *g* at 4 °C for 5 min. If needed, the lysis process was repeated. RNA was then extracted using RNeasy Plus Mini Kit (Qiagen, Cat No. 74134). After RNA quantification the RNA was hybridized with the probes and quantified using nCounter PanCancer Immune Profiling Panel for mouse according to manufacturer protocol. Analysis and statistics were performed using Rosalind software.

The mice from trial 3 (4 groups, 4 mice per group) were inoculated with B16-N4-OVA tumor and immunized as described above. On day 15, (when PBS-treated mice reached the tumor size endpoint) all mice were sacrificed, tumor masses were harvested, tumors were fixed in 4% PFA for 24 h, and then subjected to Hematoxylin and eosin staining (Histoserv, Inc.).

## ACKNOWLEDGEMENTS

We thank members of the K.S.R. lab and John Schiller for suggestions and comments on the manuscript. This work was funded by the Intramural Research Program of the National Institutes of Health, National Cancer Institute, Center for Cancer Research (C.C.R.), a C.C.R. FLEX Synergy Award (to K.S.R., K.T., and D.J.F.), and the Korean Biomedical Scientist Fellowship Program (to M.K.).

## AUTHOR CONTRIBUTIONS

Conceptualization, D.D., M.K., D.J.F., K.T., and K.S.R. Methodology, D.D., E.T., M.K., K.T., S.M.H., D.J.F., and K.S.R. Investigation, D.D., E.T., F.M., M.K., C.-H. Tai, A.M., and D.G. Formal analysis, S.M.H. Writing-Original draft, D.D. and K.S.R. Supervision, K.T., D.J.F., and K.S.R. Funding acquisition, M.K., K.T., D.J.F., and K.S.R.

## DECLARATION OF INTERESTS

K.S.R. is an inventor on a patent describing SSHEL technology that has been assigned to the U.S. government.

---

**Figure S1.**
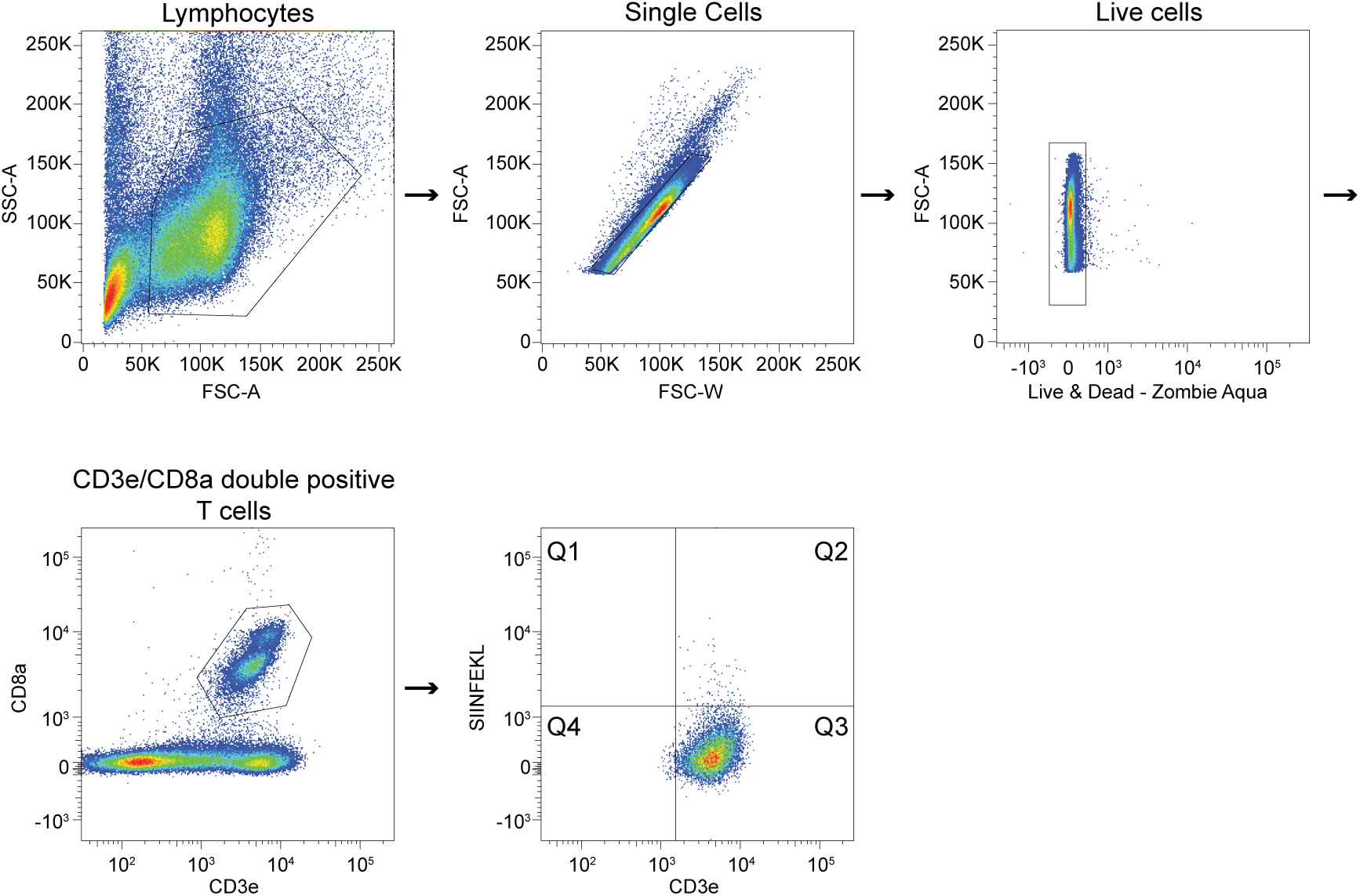
Gating strategy to isolate CD3+/CD8+ cytotoxic T cells that recognize the SIINFEKL epitope, using the H-2Kb tetramer. Related to. Fig. 4.

**Figure S2.**
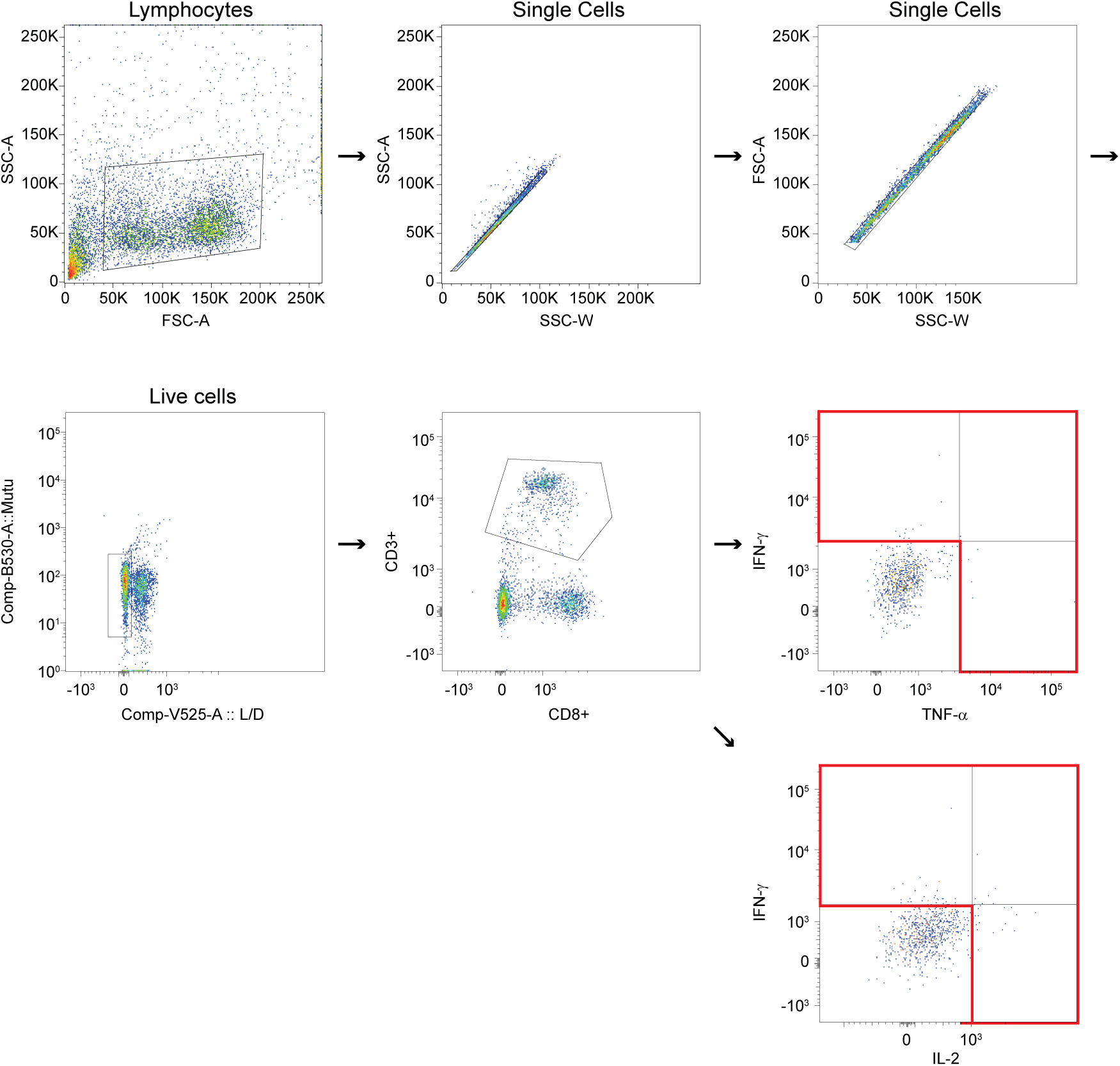
Gating strategy to isolate cytotoxic T cells releasing the indicated cytokines. Related to Fig. 5.

**Figure S3.**
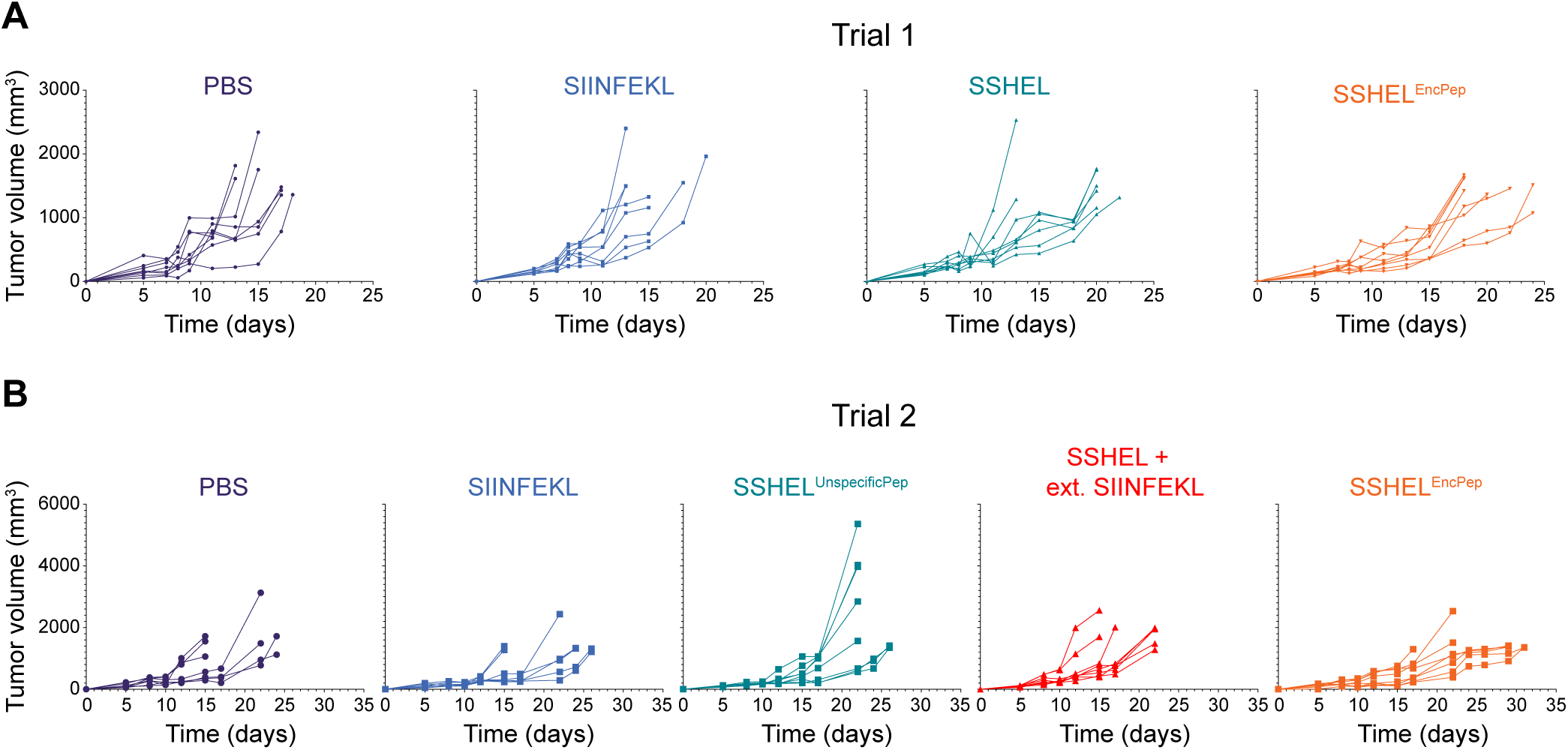
Mean weight of mice in immunization trials. (A) Trial 1 and (B) Trial 2. Data points represent mean; errors are S.D. Related to Fig. 6.

**Figure S4.**
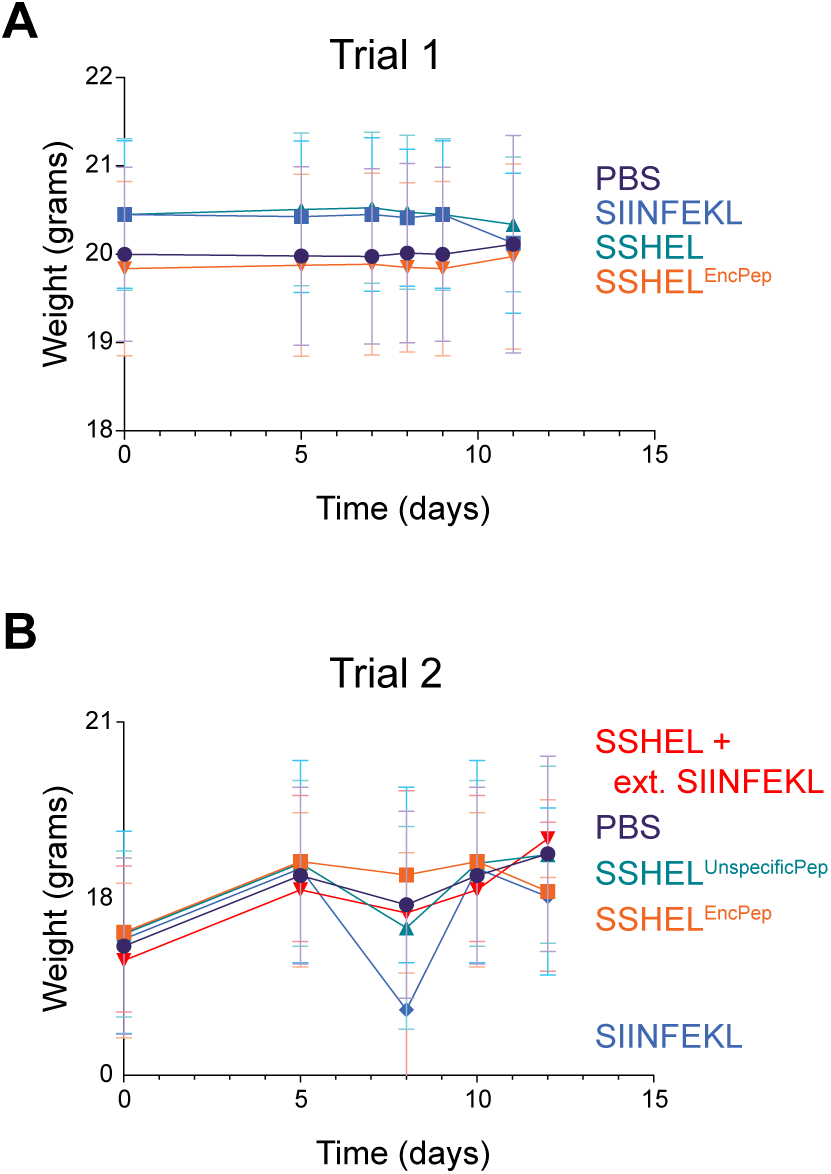
Individual tumor sizes of mice in immunization trials. (A) Trial 1 and (B) Trial 2. Treatment groups are indicated above the graphs.

**Figure S5.**
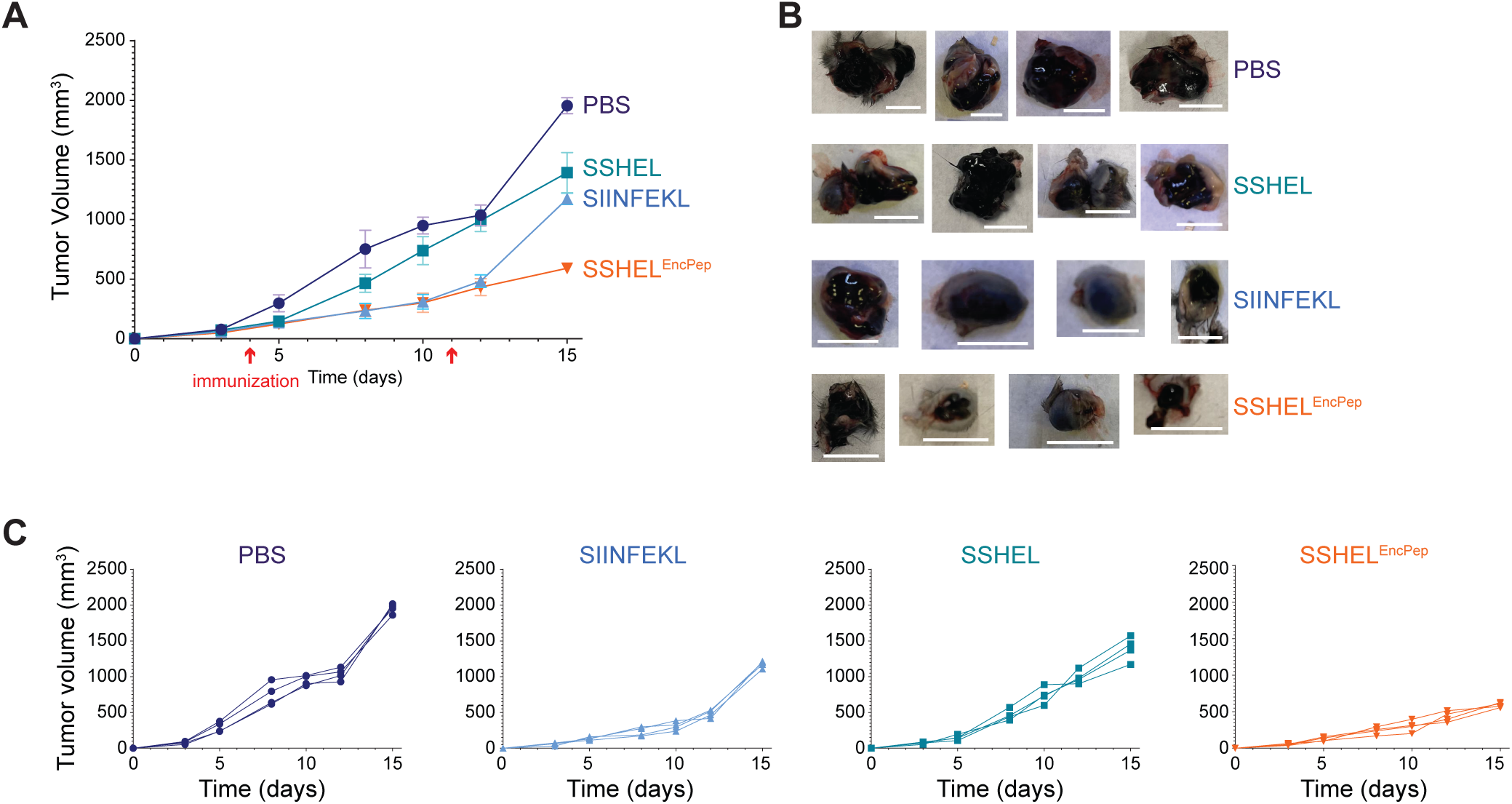
Trial 3 to assess tumor pathology. (A) Subcutaneous B16-N4-OVA tumor was introduced in C57Bl/6 mice (4 mice/treatment group). When tumors reached ∼100 mm^3^, the first inoculation was administered (7 µg peptide), followed by a second inoculation two weeks later, and tumor size was periodically measured. Mice were treated with PBS (purple), SIINFEKL peptide alone (blue), SSHEL alone (green), or SSHEL^EncPep^ (orange). (B) Images of resected tumors. Size bars: 10 mm. (C) Individual tumor sizes of each mouse.

